## Supplementary figures and images for "Placozoan secretory cell types implicated in feeding, innate immunity and regulation of behavior"

### S1 Fig

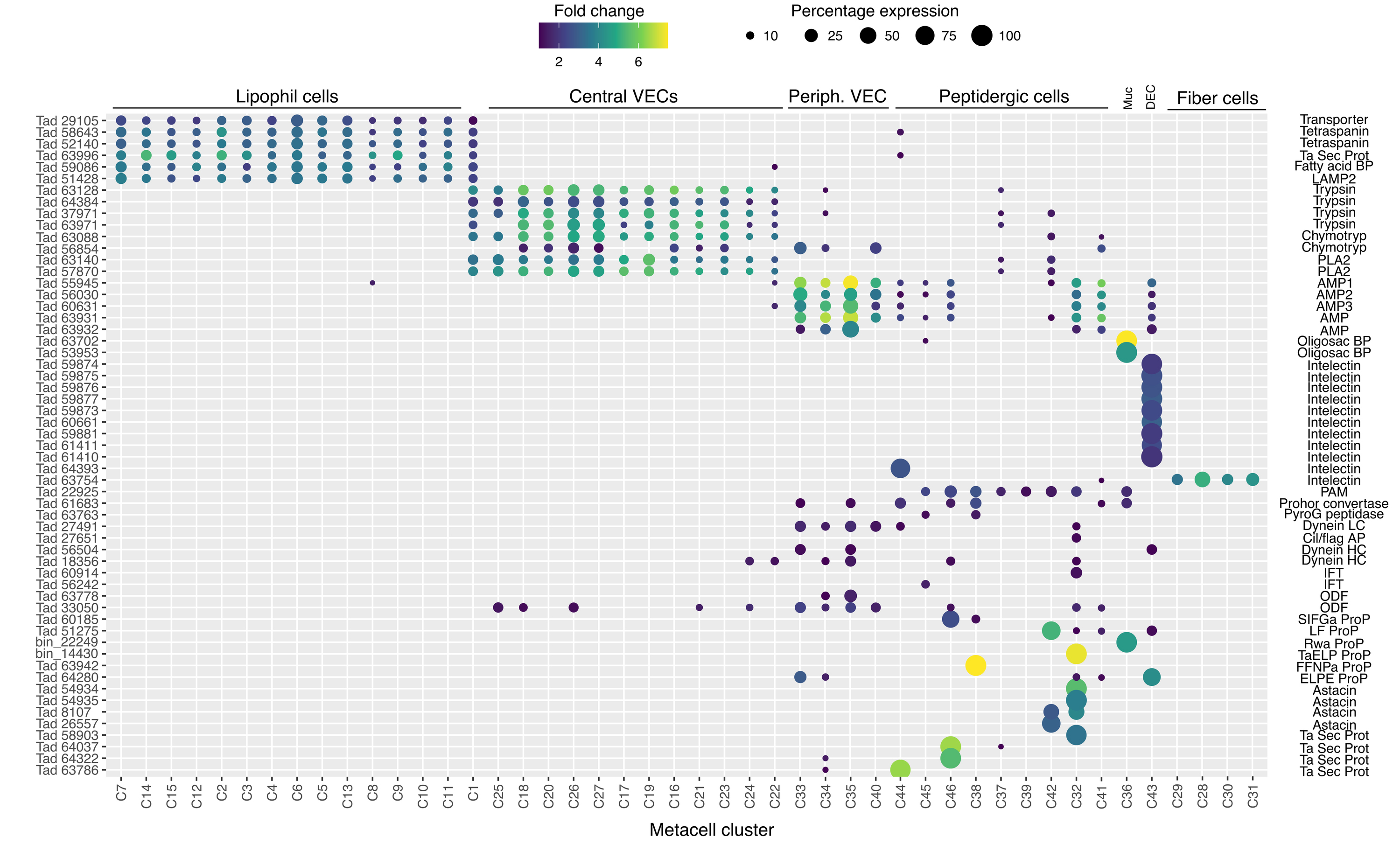

### S2 Fig

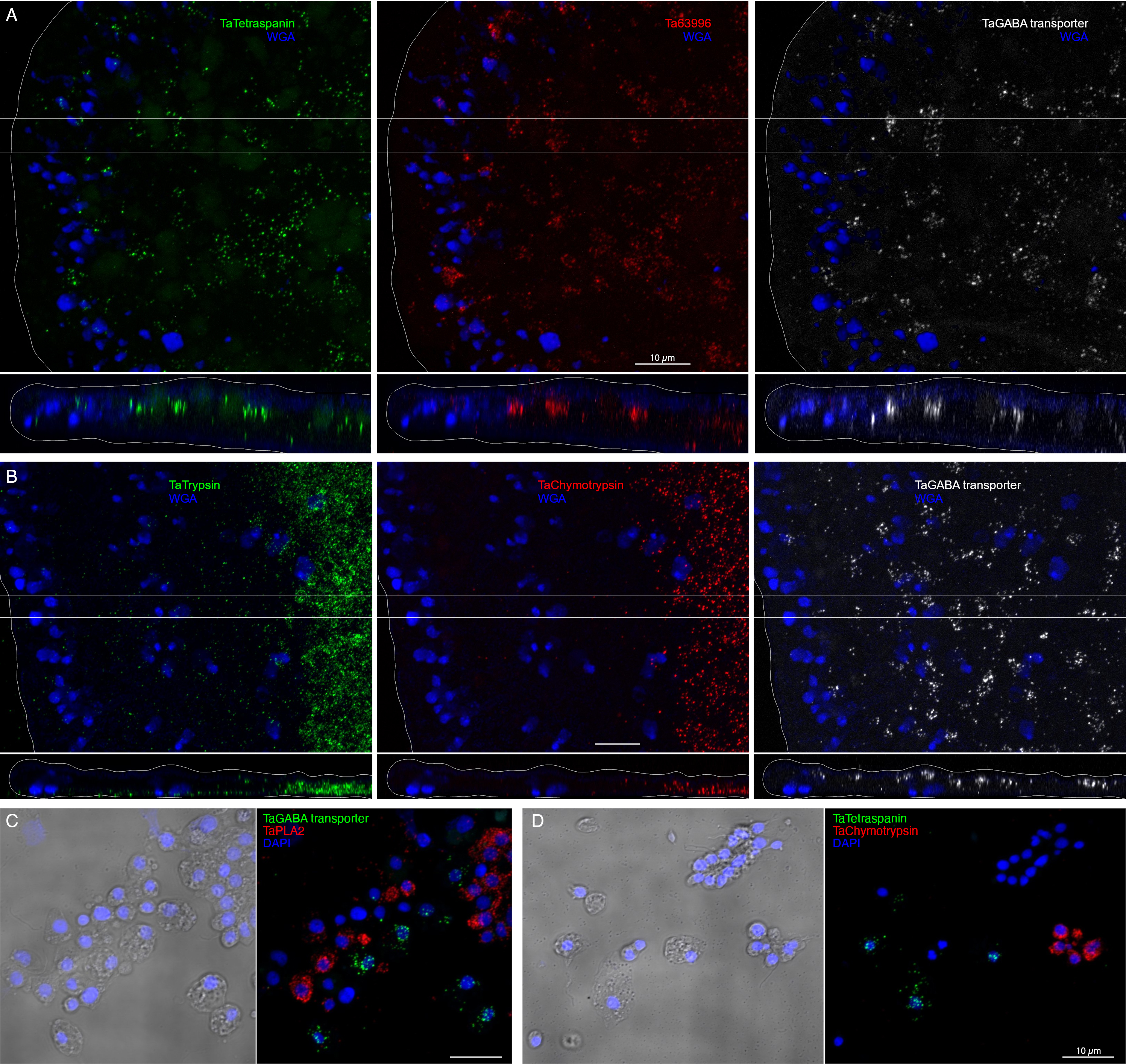

### S3 Fig

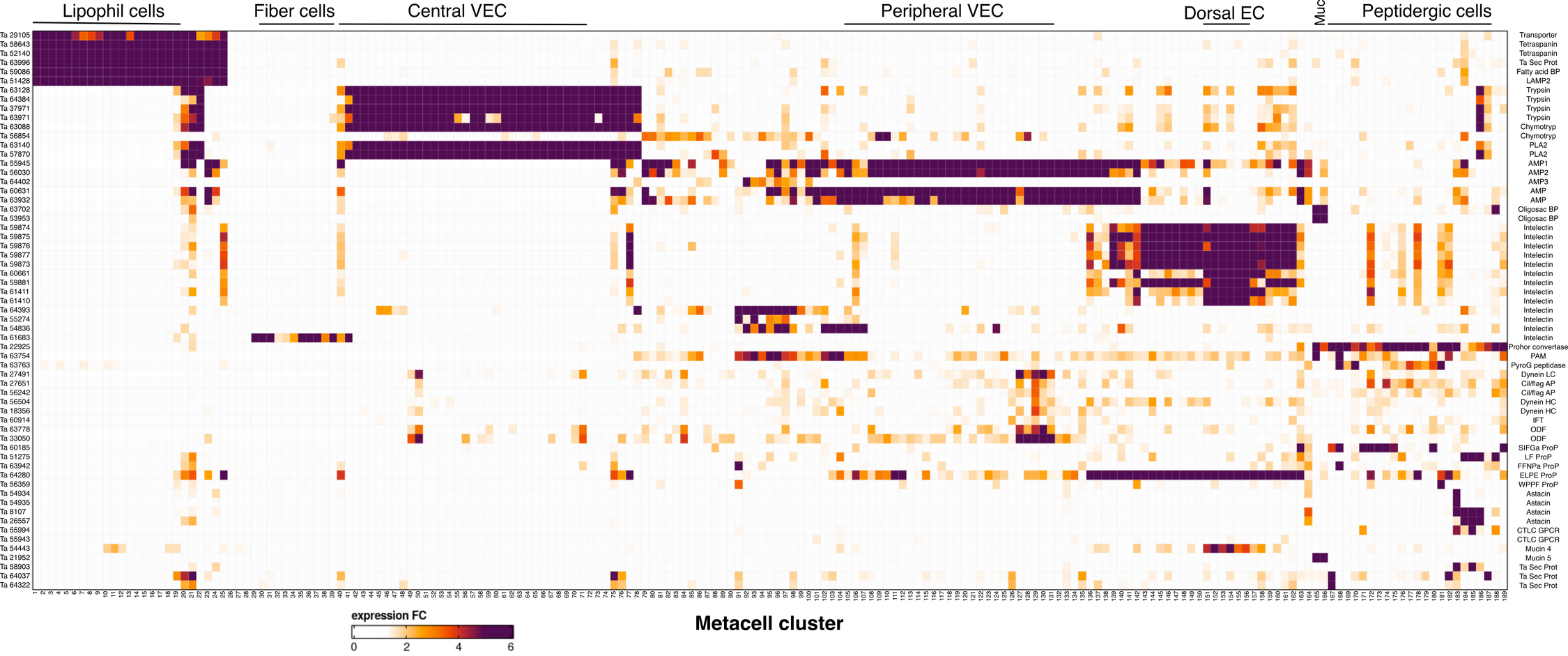

### S4 Fig

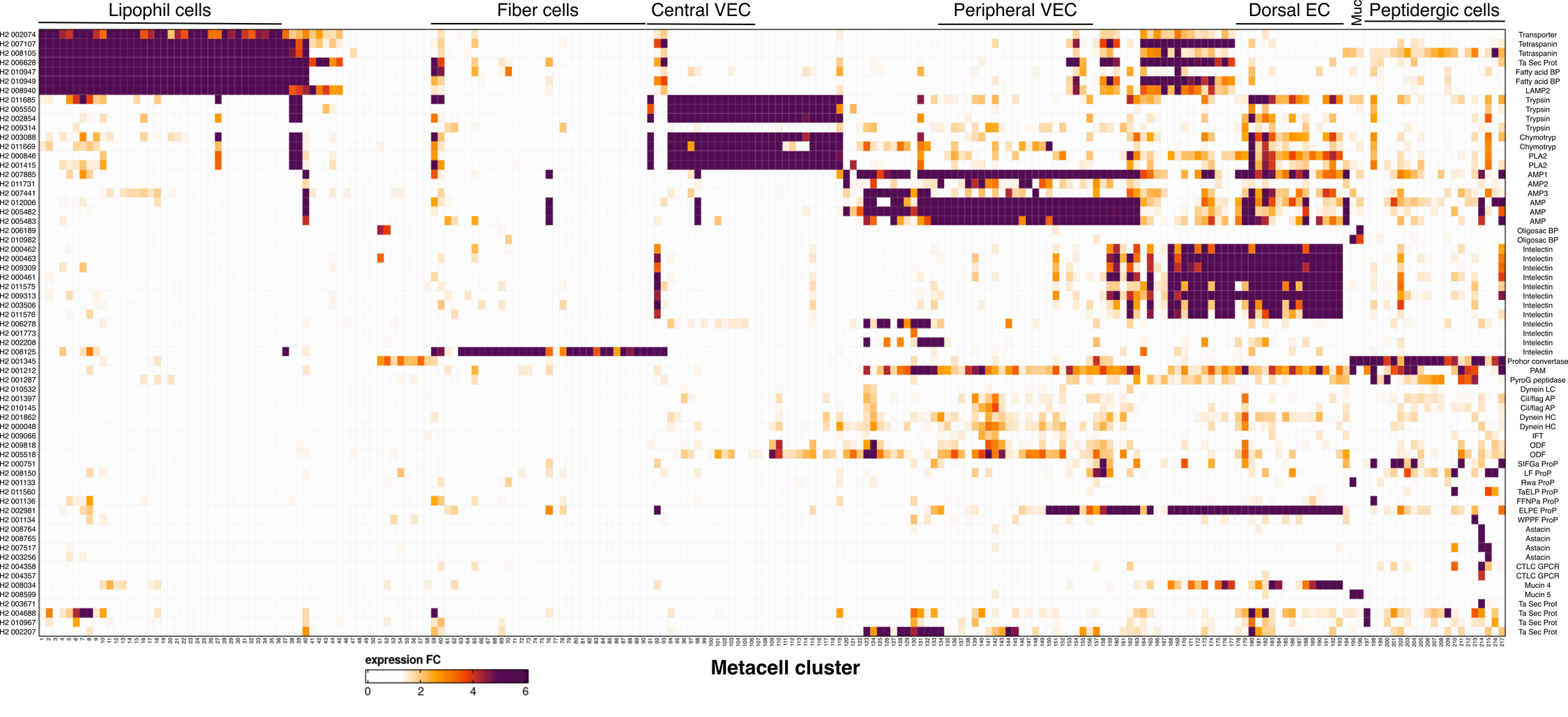

### S5 Fig

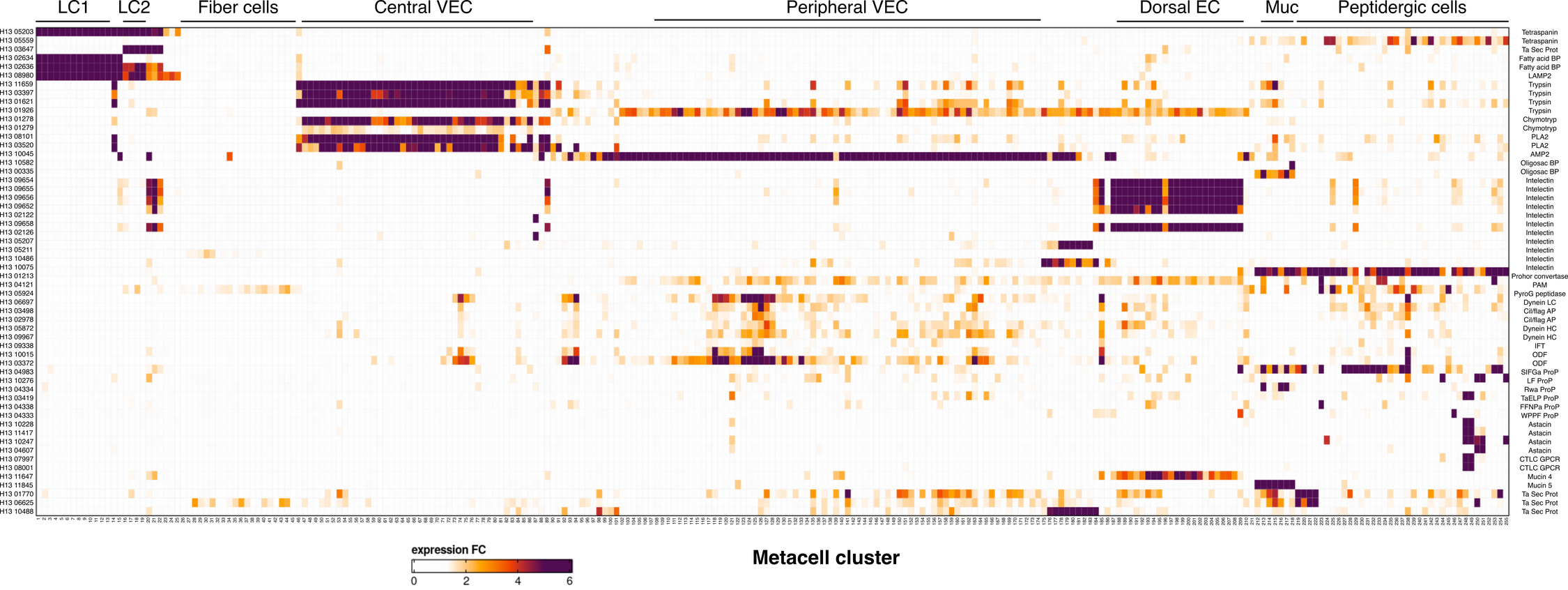

### S6 Fig

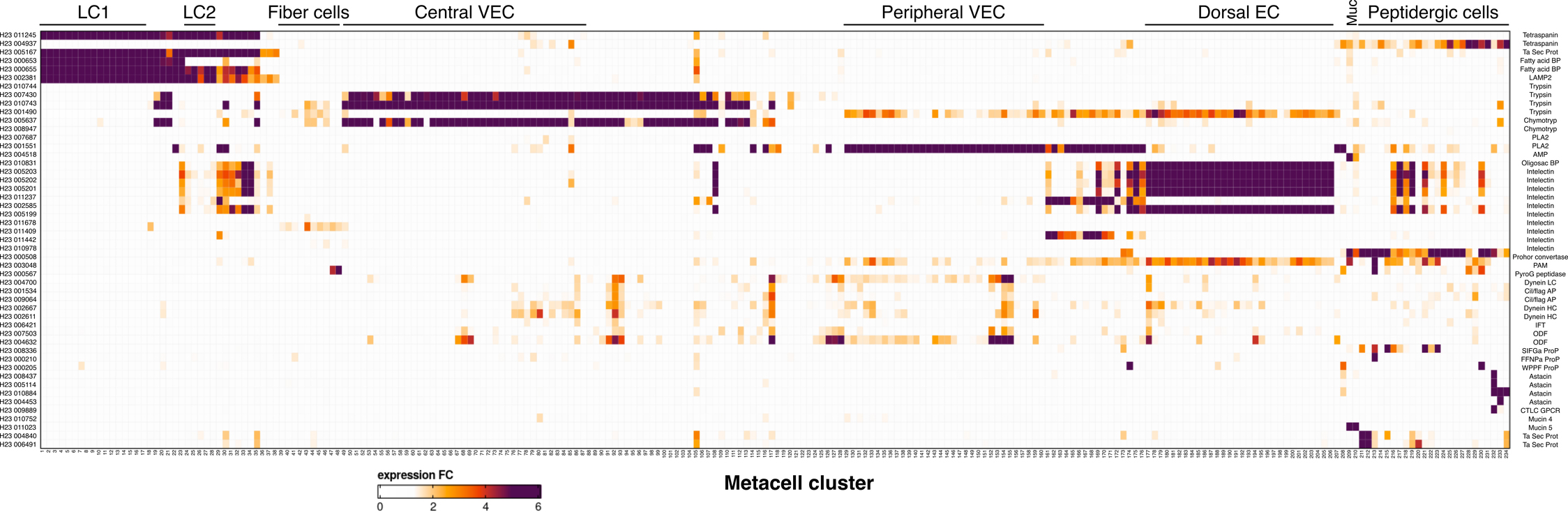

### S7 Fig

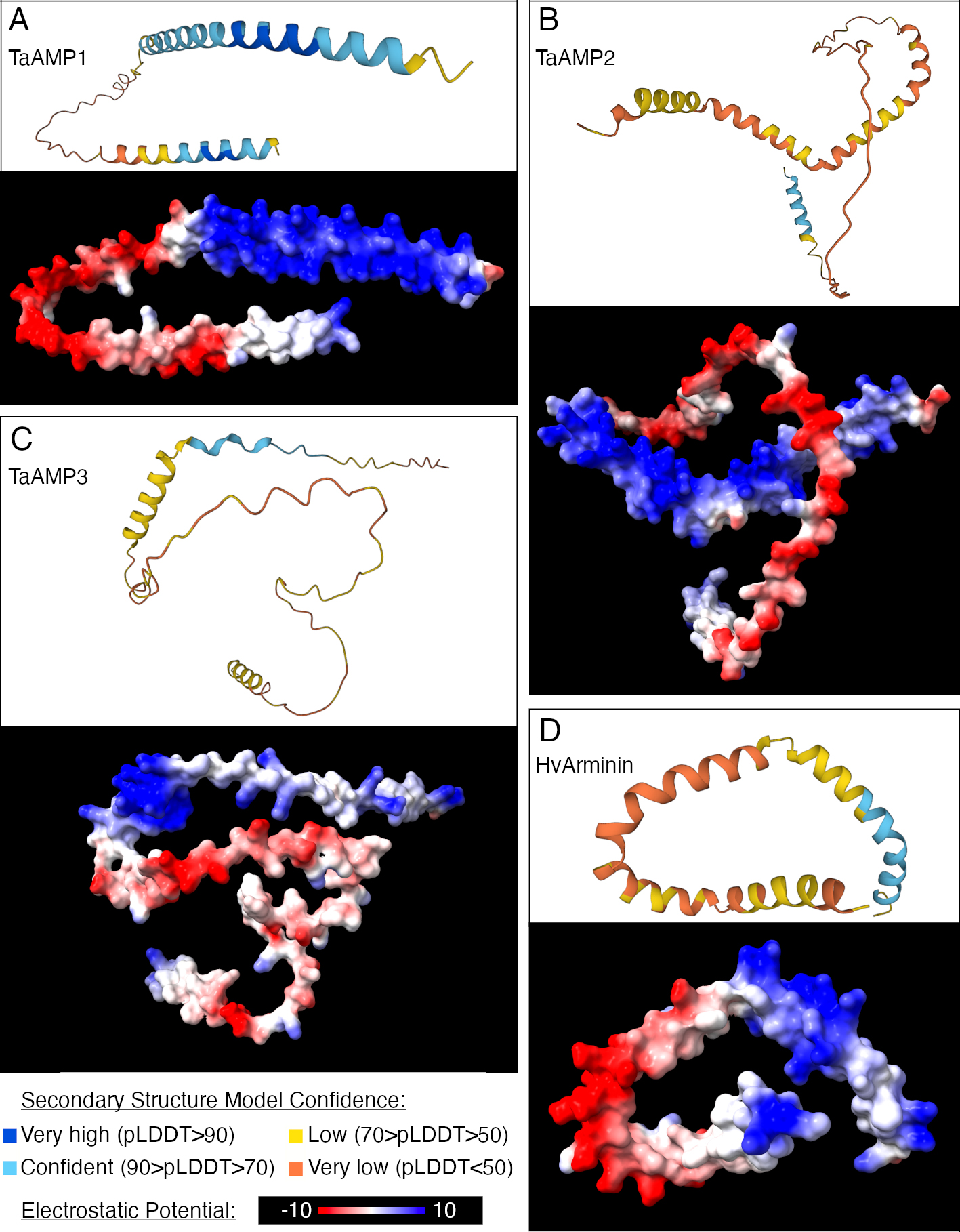

### S9 Fig

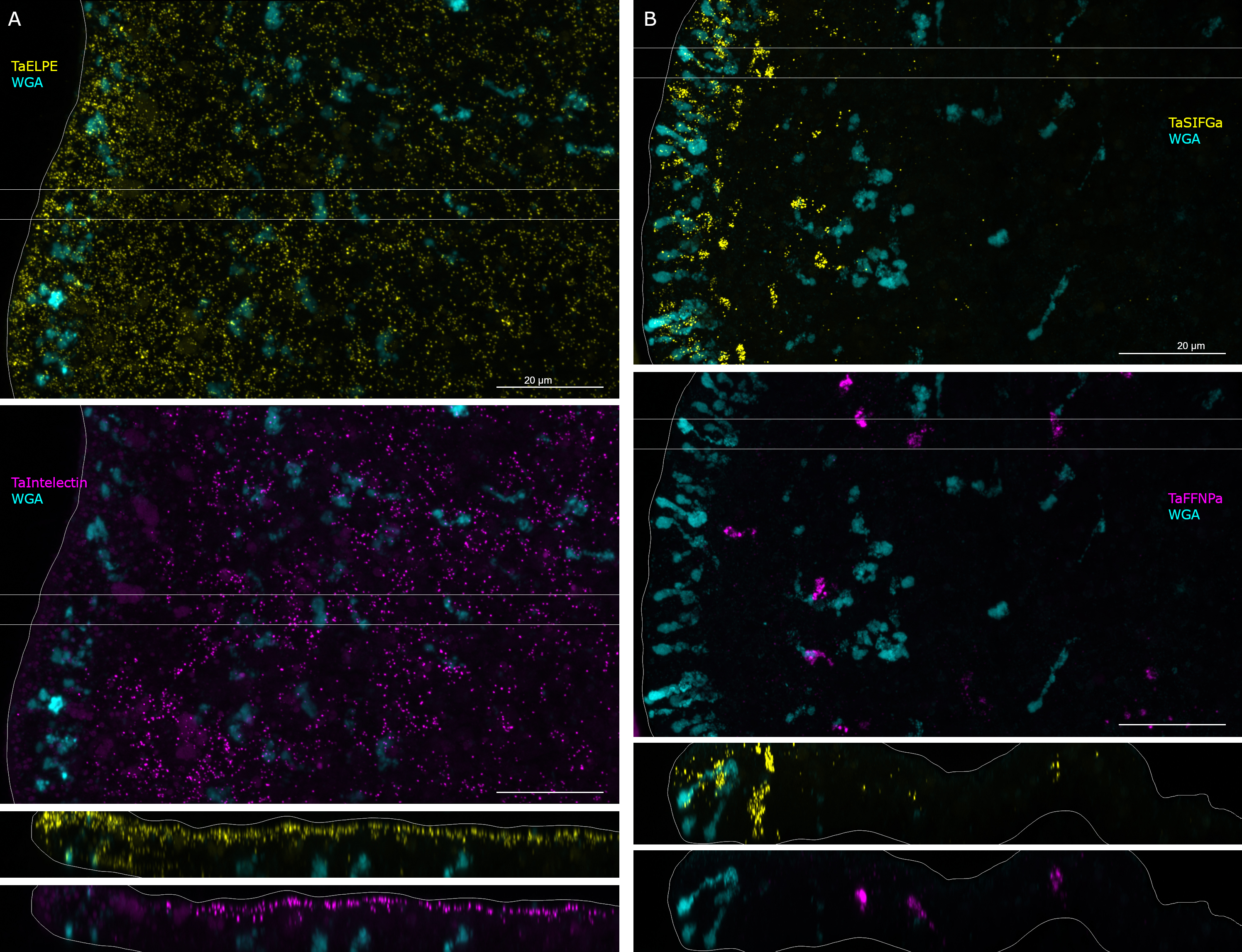

### S10 Fig

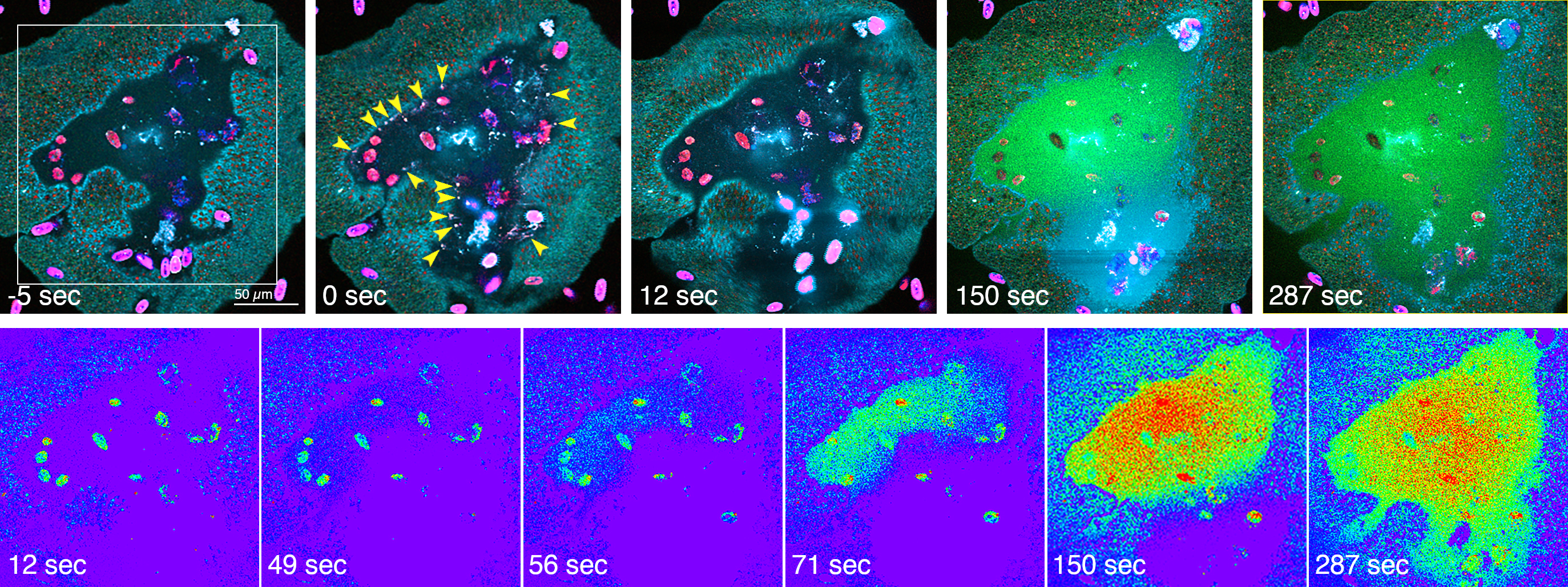
