## Supplementary material for "Placozoan secretory cell types implicated in feeding, innate immunity and regulation of behavior": S8 Text

Peptide region

Amidation

Pyro-glutamination

Cleavage site

>TH1 ELPE prepropeptide

MRSIILICLLFLFAAKVNSESFDSDDKRDLFSDEHQRNNDENVVEDGASISRQFASENDEDNNEDQIPPLGKSFELPEHRRGKSFEFPEHRRGKSFEFPERRRGKSFELPERRRGKSFELPERRRGKSFELPERRRGKSFELPERRRGKSFEFPEHRRGKSFEFPLNVLFQFGNLFRDVLARREGEIKQ

>TH2 ELPE prepropeptide

MRSIILICLLFLFAAKVNSESFDSDDKRDLFSDEHQRNNDENVVEDGASISRQFASENDEDNNEDQIPPLGKSFELPERRRGKSFEFPEHRRGKSFEFPEHRRGKSFELPERRRGKSFELPERRRGKSFELPERRRGKSFELPERRRGKSFEFPEHRRGKSFEFPVRTQTLEIIKLG

>TH1 LF prepropeptide

MRTILVFTLLVVAVSCRAISKNTDEKSKKPKKTEPKLMIGYPLFKKEDLDSQGYALFRKDDSQGYPLFRKDDSQGYPLFRKDDSQDGYALFRKDDSQPGHALFRKDDSQDGYALFRKDAQNGNSILYGHPLFKKEDQDGELSEKADTPLFKKEDSQSADSKKPIIIWKRDGPSSDSEIPMILFKKRQDDDSEKSEAKNVVSWFSQRDTRKQGFIPFKRGHKRLSYIPNSNPFKKIFLGDLSSRSEKMA*

>TH2 LF prepropeptide

MRTILVFTLLVVAVSCRAISKNTDDDTQETTKMETEPKPKLMIGYPLFKKEDLDSQDGYALFRKDDSQPGHALFRKDDSQDGYALFRKDARSENPSNIGHPLFKKEAQNGNSILYGNPLFKKEDQDGELSEKADTPLFKKEDSQSADSKRPIIIWKRDGPSSDSEIPMILFKKRQDDDSEKSEAKNVVSWFSQRDTEKQGFIPFKRGQKRLSYIPNSNPFKKIFLGDLSSRSEKMA*

>HH13 LF-1 prepropeptide

MRTLLIFVLLAIACALVNCRALEDESESWMAKRDYLLARDTKNKKKKTKLNTAAKIGFLLFKKADKLAERDFEDSLFRKSDQNSEETPAVLENIPIFRKSDRNPPDLLLFKKSDQTGGNNLFDPFKRRGIIQHGGYPWNG*

>HH13 LF-2 prepropeptide

MNKINLVTLYIIVAYAIIASSQARNVKWSRSTGTNTHNRESKLPTTFWNGNPSGTGFALFKKADDSKHLNEANRLRTPDGMGYAMFKKVHNNAHFMDKNKLKMPPGVGLPLFKKTQDSKSRKSQVANPPGFNLALFKKAQHDKKLKEDNYQFSDDSGIGLPLFKKAQHDKKPKEDNHQFSDDSGIGLPLFRKAQHDKK

>TH1 SIFGamide prepropeptide

MKQIALIFFLTAAIVFATVNAEGNLESIFNAKREDQANLKSIFGGKREDQANLKSIFGGKREDQANLKSIFGGKREDQANLKSIFGGKREDQANLKSIFGGKREDQANLKSIFGGKREDQANLKSIFGGKREDQANLKSIFGGKREDQANLKSIFGGKREDQANLKSIFGGKREDQANLKSIFGGKREDQANLKSIFGGKREDQANLKSIFGGKRDDQANLKSIFGGKRDDQANLKSIFGGKREDQANLKSIFGGKRDDQANLKSIFGGKREDQANLKSIFGGKREDQANLKSIFGGKREDQANLKSIFGGRREDQANLKSIFGGKREDQANLKSIFGGKREDQANLKSIFGGKREDQANLKSIFGGKREDQANLKSIFGGKREDQANLKSIFGGKREDQANLKSIFGGRREDQANLKSIFGGKREDQANLKSIFGGKREDQANLKSIFGGKREDQANLKSIFGGKREDQANLKSIFGGKREDQANLKSIFGGKREDQANLKSIFGGKREDQANLKSIFGGKREDQANLKSIFGGKRKDRANSKKKFGCKCKGRGNMKSMLGGKREDQANLKSIFDGKREDQANLKSIFGGKREDQANLKSIFGGKRGDQANLKSIYGGKREDQANLKSIYGGKREDQANLKSIFGGK

>TH2 SIFGamide prepropeptide

MKQIALIFFLTAAIVFATVNAEGNLESISNAKREDQANLKSIFGGKREDQANLKSIFGGKREDQANLKSIFGGKREDQANLKSIFGGKREDQANLKSIFGGKREDQANLKSIFGGKREDQANLKSIFGGKREDQANLKSIFGGKREDQANLKSIFGGKREDQANLKSIFGGKREDQANLKSIFGGKRENQANLKSIFGGKREDQANLKSIFGGKRDDQANLKSIFGGKRDDQANLKSIFGGKRDDQANLKSIFGGKREDQANLKSIFGGKREDQANLKSIFGGKREDQANLKSIFGGKREDQANLKSIFGGRREDQANLKSIFGGKREDQANLKSIFGGKREDQANLKSIFGGRREDQANLKSIFGGKREDQANLKSIFGGKREDQANLKSIFGGRREDQANLKSIFGGKREDQANLKSIFGGKRGDQANLKSIFGGKREDQANLKSIFGGKREDQANLKSIFGGKREDQANLKSIFGGMREDRGNVKSMFVGKREDQANLKSIFGGKREDQANLKSIFGGKREDQANLKSIFGGKRGDQANLKSIFGGKRADQANLKSIFGGK

>HH13 TVWGamide prepropeptide

MKSINIIFLTAAILLVSVSAGRRHDDLHKKEDTVWGGRRSDDSQRTGANLQTVWGGRRSDDSQRTGANLQTVWGGRRSDDSQRTGANLQTVWGGRRSDDSQRTGANLQTVWGGRRSDDSQRTGANLQTVWGGRRSDDSQRTGANLQTVWGGRRSDDSQRTGANLQTVWGGRRSDDDSQRTGANLQTVWGGRRSDDSQRTGANLQTVWGGRRSDDDSQRTGANLQTVWGGRRSDDSQRTGANLQTVWGGRRSDDSQRTGANLQTVWGGRRSDDDSQRTGANLQTVWGGRRSDDSQRTGANLQTVWGGRRSDDSQRTGANLQTVWGGRRSDDSKGLVPTYKLYGVDEEAMIHKRQEPTYKLYGVDEEAMIHKRQEPTYKLYGVDEEAMIS

>CH23 SFFGamide prepropeptide

MKSIYIIFFAATIVFASVNADDENVKDYFNERDNDFDVEALYEYDSKRDDDLQKKHGVNLKTFFGGKREDLQRKTGVNLKTFFGGKRDDDLQKKHGVNLKTFFGGKRDDELRKEHSVNLKNFFGREYGDLTPEVQAKTRDLTEALVALVKEIEKFVASRPNLKSFFGGKRDDADDLQRKTGVNLKTFFGGKRDDLQKKTGVNLKTFFGGKRDDLQKKTGVNLKTFFGGKRDDLQKKTGVNLKTFFGGKRDDLQRKTGVNLKTFFGGKRDDLQK

>TH1 endomorphin-like peptide prepropeptide

MDHKIKILALIVIAVAGLSSGKSMDKNGRNSVSLWTSAARDSKLAERNDQRKGYIYWETKRDENPESLALFKRKDNLLEDYPFFGNKKRQDYPFFGNKKRQDYPFFGSRKRQNLREDKVDSSDDMWDFLERDIIPFWKRNRLASIKRSRMN

>TH2 endomorphin-like peptide prepropeptide

MDHKIKILALIVIAIAGLSSGKSMDKNGRNSVSLWTSAARDSKLAERNDQRKGYIYWETKRDENPESLALFKRKDNLLEDYPFFGNKKRQDYPFFGNKKRQDYPFFGSRKRQNLREDKVDSSDDMWDFLERDIIPFWKRNRLASIKRSRMN

>HH13 endomorphin-like peptide prepropeptide

MIHKIIIVALLVIAVTDLSAGKSMDGKKDEKTLSLWTSSLGSSKASRRNDQRNGYIYWETKRDNLPFFKRRNGYPFFGGKRESDYPFFGGKREVNIINFDYTYLYRAELWDMMKREGYPYWRRDRVAAYLRSKMI

>Ta-H1 FFNPamide prepropeptide

MKTLFILLVASVALPLIIAAKDESDSKAETNKRQFNPFFKKEAEVVITNSSVKLDASKAVKVARSEDNLQKKDDQFFNPGKRDDQFFNPDKRDDQFFNPGKRDDQFFNPGKRDDQFFNPGKRDDQFFNPGKRDDQFFNPGKRDGQFFNPGKRDGQFFNPGKRDGQFFNPGKRDGQFFNPGKRDDQFFNPGRRDDQFFHSRKYDGQFFNPGKREGQFFDKGKRDDQFFNPGKRDGQFFNPGKRDAQFFNPGRRYDTQFFSPDRRRDTQFFGQRSGKDEQFFGSRGDAQFFGSRRDGQFFNPGKRDAQFFGSRDDGQFFGSKKDDQFFGHKKEDDQFFGNKKDDAQFFRNNAEETPSYYSIPRAEFMHENSGTTNNDGNNCTCDGSAPVNPFFMY

>Ta-H2 FFNPamide prepropeptide

MKTLFILLVASVALPLIIAAKDESDSKAETNKRQFNPFFKKEAEDASKAVKVARSEDNLQKKDDQFFNPGKRDDQFFNPDKRDDQFFNPGKRDDQFFNPGKRDDQFFNPGKRDDQFFNPGKRDDQFFNPGKRDGQFFNPGKRDGQFFNPGKRDGQFFNPGKRDGQFFNPGKRDDQFFNPGRRDGQFFNPGKRDDQFFHSRKYDNQFFNPGKRDGQFFNPGKREGQFFDKGKRDDQFFNPGKRDGQFFNPGKRDAQFFNPGRRYDTQFFSPDRRRDTQFFGQRSGKADEQFFGSRGDAQFFGSRRDGQFFNPGKRDAQFFGSRDDGQFFGSKKDDQFFGHKKEDDQFFGNKKDDAQFFRNNAEETPSYYSIPRAEFMHENSGTTNNDGNNCTCDGSAPVNPFFVY

>Hoilungia-H13 FFGQamide prepropeptide

MKILFIFLIASMALPVIVSAKDESNEKSEINRRQFNPFFKKEVKVKIIIILFLIVSLLINLDDQFFGQGKRDAQFFGQGKRDDQFFGQGKRDDQFFGQGKRDDQFFGQGKRDDQFFGQGKRDDQFFGQGKREDQFFGQGKRDDQFFGQGKRDAQFFGQGKRDDQFFGQGKRNDQFFGQGKRDAQFFGQGKRDNQFFGQGKRDAQFFGQGKRDAQFFGQGKRDDQFFGQGKRDDQFFGQGKRDDQFFGQGKRDDQFFGQGKRDNQFFGQGKRDGQFFGNRHANKQFFGNGRGKFCLCFNMLHFMAYAVYV

>Cladtertia-H23 FFGQamide prepropeptide

MKILFIFLIASMALPAIISAKDEVNDKSEINRRQFNPFFKKEAKDTPKAAKAERSGMSTNFTINITSENLKQFSIILFNDHLMINFLAKANAMINFLDKANVVRIDFILVFIVKLRLKYSIFYSTDDQFFGQGKRDDQFFGQGKRDAQFFGQGKRDDQFFGQGKRDAQFFGQGKRDAQFFGQGKRDAQFFGQGKRDNQFFGQGKRDAQFFGQGKRDGQFFGNGRADKQFFGNGRDTQFFGNGRADTQFFGNGRDTQFFGNGRDTQFFGNRGDTQFFNPDRRDDAQFFGNRGDYQFFGNRDDGQFFGHKKDDQFFGSRKGI

PIIALLKQ

>Ta-H1 RWamide prepropeptide

MLTNRFIIWILFLGITTAQNVAKGKAQIGNHKSVFLKNEATRPERDQPPRWGRDQPTRWGRDQPPRWGRDQPPRWGRDQPSRWGRDQPPRWGRDQPPRWGRDQPPRWGRDQPPRWGRDQPPRWGRDQPPRWGRDQPPRWGRDQPPRWGRDQPPRWGRDQPPRWGRDQPPRWGRDQPPRWGGDQLPEMEKNHAPPRWGRDQYSWWNQEQYPSRWGREYSTPDNTAEKLLDSLTHQSENAKKNNFQEINSDSNSGNESAVHRLFSNKLKNQKAKSDSNKLMNSFSGSESISRPREKSLKRSETLDNMRIDLI

>Ta-H2 RWamide prepropeptide

MLTNRFIIWILFLGITTAQNVAKGKAQIGNHKSVFLKNEATRPERDQPPRWGRDQPTRWGRDQPPRWGRDQPPRWGRDQPSRWGRDQPPRWGRDQPPRWGRDQPPRWGRDQPPRWGRDQPPRWGRDQPPRWGRDQPPRWGRDQPPRWGRDQPPRWGGDQLPEIEKNYAPPRWGRDQYSWWNQEQYPSRWGREYSTPDNTAEKLLDSLTHQSENAKKNNFQEINSDSNSGNESAVHRLFSNKLKNQKAKSDSNKLMNSFSGSESISRPREKSLKRSETLDNMRIDLI

>Hoilungia-H13 RWamide prepropeptide

MLTNRLIILLLLGIATAKNVVKDNTAADVSDHSRFSKDQTYIIKSDQPPRWGRDQPPRWGRNQPPRWGRNQPLSWEYDQSLIYERDQPPRWGRDQPPRWGRNQPPRWGRDQPPRWGRNQPPRWGRDQPPRWGRDQPPRWGRDQPPRWGRDQPPRWGRDQPPRWGRDQPPRWGRDQPPRWGRNQPMELQVDHAPPRWGREQFSWWNEDKYPNRWGRKHHSSADNAKEESLDILMSQSKNTLDNMHKVIGTDNDAIVIGGLPSSINQANQDDKAATKTNDMTENLSVTE

>Cladtertia-H23 RWamide prepropeptide

MKMLANRLIILLLLGITTAQNVVKDKTVASIRDHSKLSKDTPYVVKRDQPPRWGRDQPPRWGRDQPPRWGRDQPPRWGRDQPPRWGRDQPPRWGRHQPPRWGRDQPPRWGRDQPPRWGRDQPPRWGRNQPPRWGRDDQPPRWGRDDQPPRWGRDDQPPRWGRDDQPPRWGRDDQPPRWGRDDQPPRWGRDDQPPRWGRDQPPRWGRSQPVELEDNQAPPRWGRDQFSWWNKDKYPNRWGRENHSSADKIRDESLDILTHQSKNMVDNIQKIIDGTNNDATIVDRLLSSKNQDDKATTKTNNVAEKLSLSE

>Ta-H1 WPPF prepropeptide

MYRLSLCCIIILVLFANEIEPKFAKPKEDIPWNLQRRSNANNLKSRSDSAKLSNTEHKKKDLVAEEQSHPIFGKGLVNEAKKTSRNEALQYNGWPPFRREDESKQYNGWPPFRREDESKQYNGWPPFRRSDELTQYNGWPPFRRNDGKEQYNGWPPFRRNAGMMQYNGWPPFRRDDEKMQYNGWPPFRREDREKQYNGWPPFRRDDEVMQYNGWPPFRRSEAVQYNGWPPFRRDDQQNKPYNGWPPFRREDQQNKPYNGWPPFRRDDQQNKPYNGWPPFRRNDQQKKPYNGWPPFRRNN

>Ta-H2 WPPF prepropeptide

MYRLSLCCIIILVLFANEIEPKFAKPKEDIPWNLQRRSNANNLKSRSDSAKLSNTEHKKKDLVAEEQSHPIFGKGLVNEAKKTSRNEALQYNGWPPFRREDESKQYNGWPPFRREDESKQYNGWPPFRRSDELTQYNGWPPFRRNDGKEQYNGWPPFRRNAGMMQYNGWPPFRRDDEKMQYNGWPPFRRDDDGKQYNGWPPFRRDDEVMQYNGWPPFRRSEAVQYNGWPPFRRDDQQNKPYNGWPPFRREDQQNKPYNGWPPFRRDDQQNKPYNGWPPFRRNDQQKKPYNGWPPFRRNN

>Hoilungia-H13 WPPF prepropeptide

MFRLPIYLTIVLVLCAHQIEPKHIKSKNDLLWNLQRRSHANLKAHSDLSKLANTEQKKKDSAAKEQTHPIFGKGDVKEAKASRDEISQYNGWPPFRRDNSKQYNGWPPFRSRSDAAAEVEQYNGWPPFRSRSDEMTEVEQYNGWPPFRRDDELKQYNGWPPFRREDQQNKPYNGWPPFRRDDQQNKPYNGWPPFRRDDQQNKPYNGWPPFRRDEQQNKPYNGWPPFRAIY
